# A framework for translation of genomic responses from mouse models to human inflammatory disease contexts

**DOI:** 10.1101/346122

**Authors:** Douglas K. Brubaker, Elizabeth A. Proctor, Kevin M. Haigis, Douglas A. Lauffenburger

## Abstract

The high failure rate of therapeutics showing promise in mouse disease models to translate to patients is a pressing challenge in biomedical science. However, mouse models are a useful tool for evaluating mechanisms of disease and prioritizing novel therapeutic agents for clinical trials. Though retrospective studies have examined the fidelity of mouse models of inflammatory disease to their respective human *in vivo* conditions, approaches for prospective translation of insights from mouse models to patients remain relatively unexplored. Here, we develop a semi-supervised learning approach for prospective inference of disease-associated human *in vivo* differentially expressed genes and pathways from mouse model experiments. We examined 36 transcriptomic case studies where comparable phenotypes were available for mouse and human inflammatory diseases and assessed multiple computational approaches for inferring human *in vivo* biology from mouse model datasets. We found that a semi-supervised artificial neural network identified significantly more true human *in vivo* associations than interpreting mouse experiments directly (95% CI on F-score for mouse experiments [0.090, 0.175], neural network [0.278, 0.375], p = 0.00013). Our study shows that when prospectively evaluating biological associations in mouse studies, semi-supervised learning approaches combining mouse and human data for biological inference provides the most accurate assessment of human *in vivo* disease and therapeutic mechanisms. The task of translating insights from model systems to human disease contexts may therefore be better accomplished by the use of systems modeling driven approaches.

**Author Summary:** Comparison of genomic responses in mouse models and human disease contexts is not sufficient for addressing the challenge of prospective translation from mouse models to human disease contexts. Here, we address this challenge by developing a semi-supervised machine learning approach that combines supervised modeling of mouse experiment datasets with unsupervised modeling of human disease-context datasets to predict human *in vivo* differentially expressed genes and pathways as if the model system experiment had been run in the human cohort. A semi-supervised version of a feed forward artificial neural network was the most efficacious model for translating experimentally derived mouse molecule-phenotype associations to the human *in vivo* disease context. We find that computational generalization of signaling insights from mouse to human contexts substantially improves upon direct generalization of mouse experimental insights and argue that such approaches can facilitate more clinically impactful translation of insights from preclinical studies in model systems to patients.

## INTRODUCTION

Generalization of insights from nonhuman disease model systems to the human *in vivo* context remains a persistent challenge in biomedical science. The association of molecular features with a phenotype in model systems, particularly for inflammatory pathologies, often does not hold true in the corresponding human *in vivo* indication, due to some combination of both the fidelity of the experimental system to human *in vivo* biology and the inherent complexity of human inflammatory disorders (1-7). Though it is now possible to collect clinical samples from patients and associate molecular features with clinical phenotypes in a human *in vivo* context, there is a discrepancy between the type and quantity of phenotypes measurable in patient cohorts and those phenotypes investigable by use of mouse disease models. Outside of a clinical trial, novel perturbations to the disease system cannot be directly investigated in the patient *in vivo* context, whereas mouse disease model systems can be used to study the impact of innumerable perturbations to the disease system and to associate molecular features, such as differentially expressed genes (DEG), with these responses. As a consequence of this discrepancy, murine and other animal model systems of disease are likely to remain an important part of biomedical research. Therefore, methods for improving the generalizability of mouse-derived molecular signatures to the human *in vivo* disease context are needed to facilitate more impactful translational research.

The utility of mouse models for studying inflammatory pathologies in particular was recently assessed by a pair of studies examining the correspondence between gene expression in murine models of inflammatory pathologies and human contexts (1, 2). The human and mouse microarray cohorts assembled by the two studies had the rare property that mouse molecular and phenotype data were well matched to human *in vivo* molecular and phenotype data. This property enabled systematic examination of the similarities and discrepancies between mice and humans. These studies analyzed the same cohorts of mouse and human studies and came to conflicting conclusions about the relevance of mouse models for inflammatory disease research, with Seok *et al.* concluding that mouse models poorly mimic human pathologies, whereas Takao *et al.* concluded that mouse models usefully mimic human pathologies (1, 2). The key methodological difference between the two studies was that while Seok *et al*. examined genes changed in either mouse or human contexts, Takao *et al.* examined only those genes significantly changed in both contexts (1, 2). However, in prospective studies of disease mechanisms and therapeutic efficacy, the corresponding mouse and human *in vivo* datasets and perturbations are rarely available. The scenario described by Takao *et al.* where genes of interest can be selected on the basis of changing in both human and mouse contexts is therefore somewhat artificial since prospective investigations will have to proceed on the basis of genes changing in the mouse contexts alone. Neither Seok *et al.* nor Takao *et al.* examined approaches addressing this real-life scenario of predicting human *in vivo* biology from mouse model datasets either *a priori* or with the aid of computational techniques.

The aim of our study here is to address the challenge of prospective inference of human biology from a model system study by developing a machine learning approach for inferring human biological associations *as if* the model system study had been conducted in a human cohort. Within this framework, a machine learning approach is judged successful if it correctly predicts a higher proportion of human DEGs and enriched signaling pathways than were implicated by the corresponding mouse disease model prior to any computational analysis. The essence of our approach is to apply a machine-learning classifier to assign synthetic phenotypes derived from those in a mouse dataset to molecular datasets of disease-context associated human samples. These synthetic phenotype labels of the human samples are then used for differential expression and pathway enrichment analysis to derive a set of predicted molecule-phenotype associations for the human samples. We were able to assess the efficacy of this approach by testing it on the datasets from the Seok and Takao studies, where mouse phenotypes and gene expression data were matched to patient clinical phenotypes and gene expression data (1, 2, 8-20).

We found that while mouse experiments alone failed to capture a large portion of human *in vivo* molecule-phenotype relationships, as noted by Seok *et al.*, using these datasets to train computational models produced more accurate and comprehensive predictions of human *in vivo* biology. In particular, a semi-supervised artificial neural network identified significantly more human *in vivo* DEGs and pathways than mouse models alone or other machine learning approaches examined here. Our results suggest that computational generalization of insights from mouse model systems to human contexts effectively captures a more substantial extent of human *in vivo* disease biology and that such approaches may facilitate more clinically impactful interpretation of model system insights.

## RESULTS

### Developing a framework for mouse-to-human genomic insight translation

We assembled a cohort of mouse-to-human translation case studies from the datasets analyzed in Seok *et al.* and Takao *et al.* (1) (2). This cohort contained mouse and human datasets for two conditions where human disease etiology can be easily mimicked in the mouse (burn and trauma) and two conditions of complex, polygenic etiology difficult to mimic in mouse models (endotoxemia and sepsis) (Table 1). We defined case studies as all pairs of mouse and human datasets for the same disease condition. Datasets with multiple disease conditions, or multiple microarray platforms were separated into different case studies by disease and microarray platform. By constructing case studies in this manner, multiple mouse strains and experimental protocols could be compared to different presentations of that same disease in independent human cohorts. Our final cohort consisted of 36 mouse-to-human translation case studies in which comparable mouse and human phenotypes existed, enabling assessment of the correspondence between mouse models and humans as well as the performance of algorithmic approaches for mouse-to-human biological translation (Table 2).

**Table 1:** Cohort of mouse and human inflammatory pathology microarray datasets. Datasets are identified by Gene Expression Omnibus (GEO) accession numbers and microarray platorms. Conditions and sample sizes are shown as well as tissue source for each dataset. Inflammatory disease inductions include lipopolysaccharide (LPS), cecal ligation and puncture (CLP), streptococcus pneumoniae serotype 2 (SPS2), and staphylococcus aureus (SA) exposure.

| Species | Disease | GEO<br>Accession | Microarray<br>Platform(s) | Conditions and Sample<br>Sizes | Tissue |
| --- | --- | --- | --- | --- | --- |
| Human | Burn | GSE37069 | GPL570 | Control (n = 37)<br>Burn (n = 553) | White Blood Cells |
| Human | Trauma | GSE36809 | GPL570 | Control (n = 37)<br>Trauma (n = 216) | Whole Blood |
| Human | Endotoxemia | GSE3284 | GPL96<br>GPL 97 | Control (n = 16)<br>Endotoxin (n = 76) | Whole Blood |
| Human | Sepsis | GSE13904 | GPL570 | Control (n = 18)<br>Sepsis (n = 158) | Whole Blood |
| Human | Sepsis | GSE9960 | GPL570 | Control (n = 16)<br>Sepsis (n = 54) | Peripheral Blood<br>Mononuclear Cells |
| Human | Sepsis | GSE28750 | GPL570 | Control (n = 20)<br>Sepsis (n = 21) | Whole Blood |
| Human | Sepsis | GSE13015 | GPL6106<br>GPL6947 | Control (n = 29)<br>Sepsis (n = 77) | Whole Blood |
| Mouse<br>(C57BL/6J) | Burn | GSE7404 | GPL1261 | Control (n = 16)<br>Burn (n = 16) | Leukocytes |
| Mouse<br>(C57BL/6J) | Trauma | GSE7404 | GPL1261 | Control (n = 16)<br>Trauma (n = 16) | Leukocytes |
| Mouse<br>(C57BL/6J) | Endotoxemia:<br>LPS | GSE7404 | GPL1261 | Control (n = 8)<br>Endotoxemia (n = 8) | Leukocytes |
| Mouse<br>(C57BL/6J) | Endotoxemia:<br>LPS | GSE5663 | GPL81 | Control (n = 4)<br>Endotoxemia (n = 5) | Blood |
| Mouse<br>(C57BL/6J) | Sepsis: CLP | GSE5663 | GPL81 | Control (n = 4)<br>Mild Sepsis (n = 5)<br>Sepsis (n = 5) | Blood |
| Mouse<br>(C57BL/6J) | Sepsis: SPS2 | GSE26472 | GPL6778 | Control (n = 3)<br>Sepsis (n = 8) | Blood |
| Mouse<br>(A/J and<br>C57BL/6J) | Sepsis: SA | GSE19668 | GPL1261 | Control A/J (n = 5)<br>Control C57BL6J (n = 5)<br>Sepsis A/J (n = 20)<br>Sepsis C57BL6J (n = 20) | Blood |

**Table 2:**
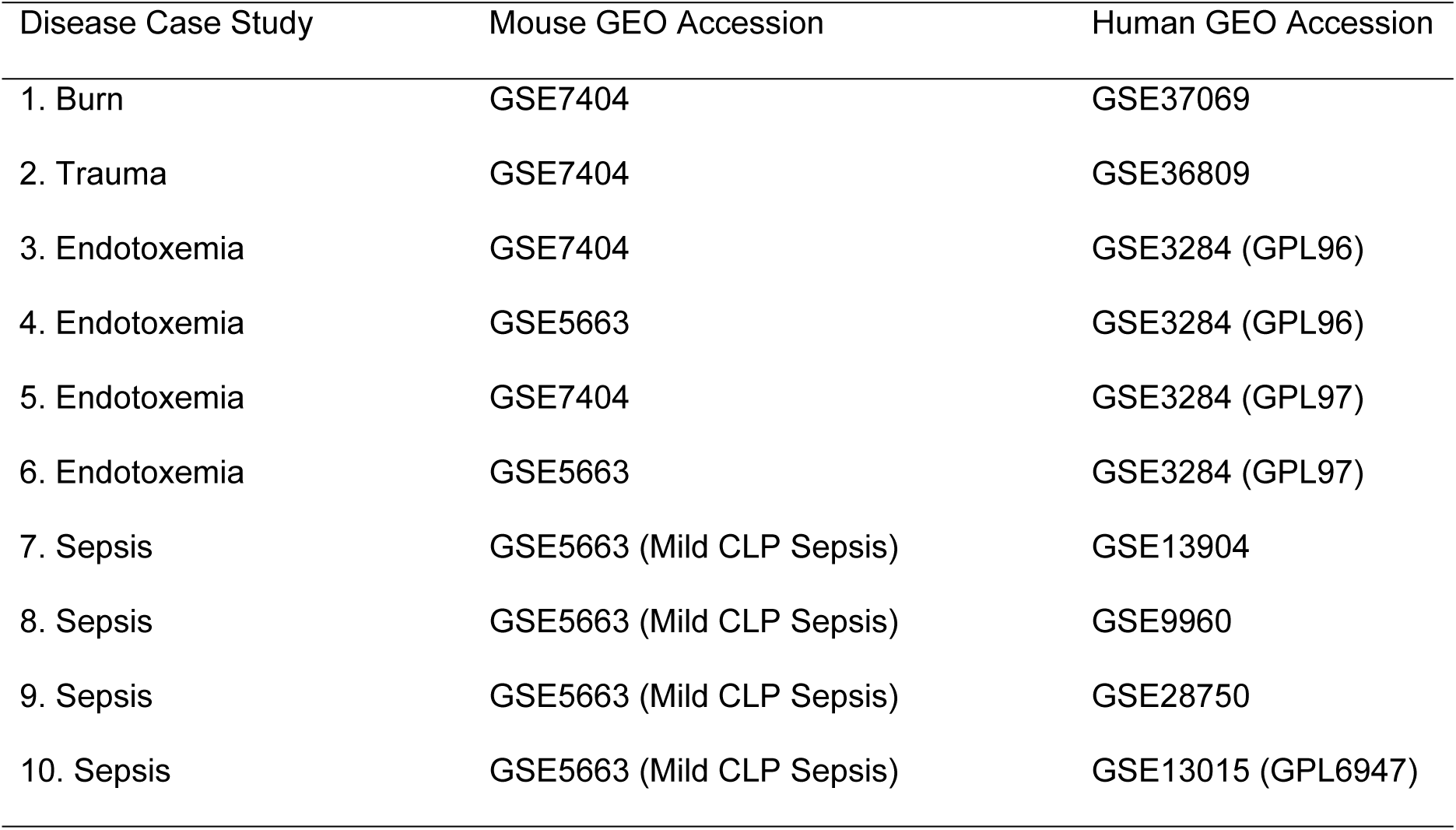
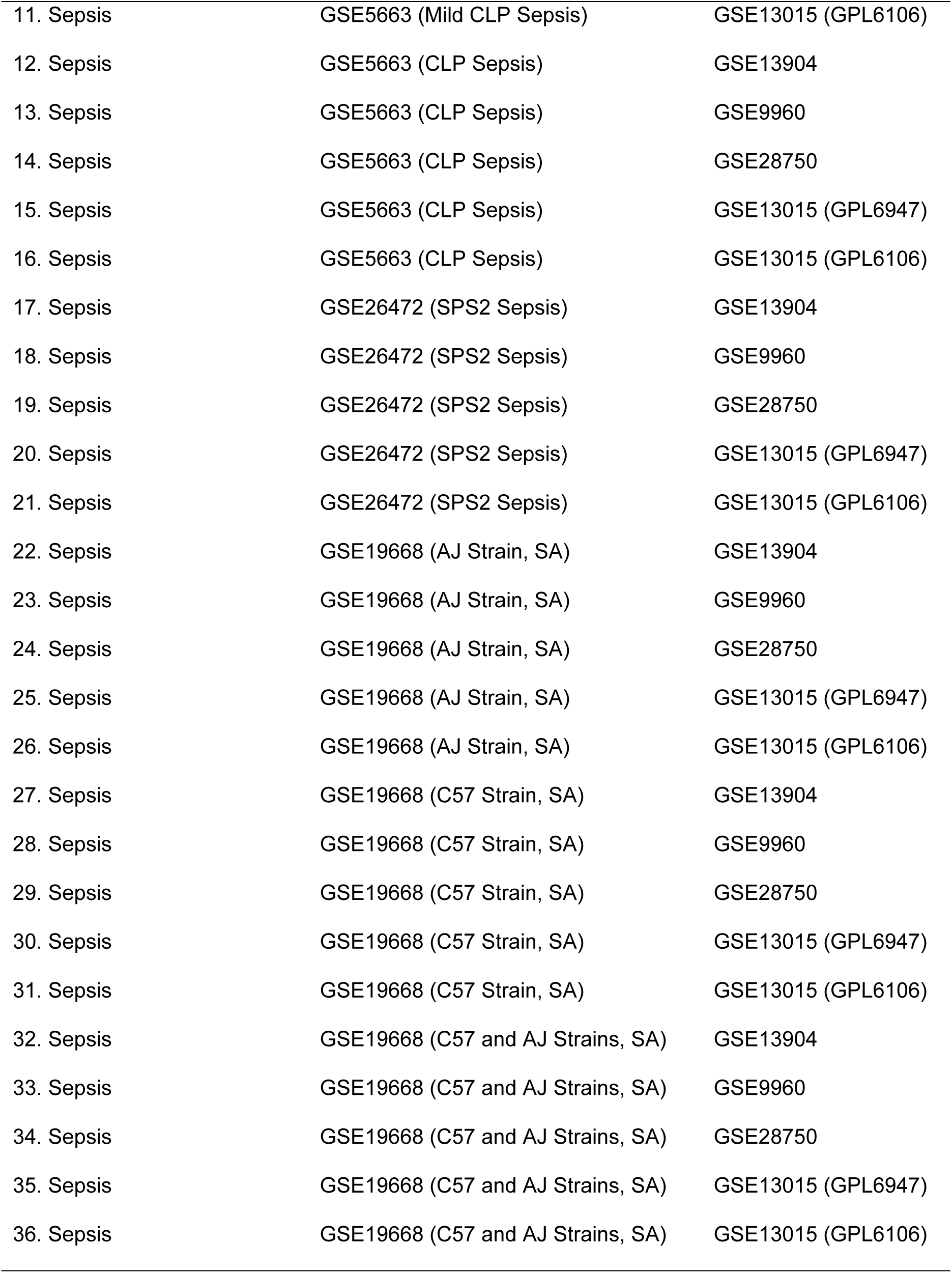
Enumeration of mouse-to-human translation case studies. Case studies are defined by all possible combinations of mouse and human datasets (GEO-GSE number) for a given mouse strain, microarray platform (GPL number), and method of disease induction when multiple methods are associated with the same dataset.

| Disease Case Study | Mouse GEO Accession | Human GEO Accession |
| --- | --- | --- |
| 1. Burn | GSE7404 | GSE37069 |
| 2. Trauma | GSE7404 | GSE36809 |
| 3. Endotoxemia | GSE7404 | GSE3284 (GPL96) |
| 4. Endotoxemia | GSE5663 | GSE3284 (GPL96) |
| 5. Endotoxemia | GSE7404 | GSE3284 (GPL97) |
| 6. Endotoxemia | GSE5663 | GSE3284 (GPL97) |
| 7. Sepsis | GSE5663 (Mild CLP Sepsis) | GSE13904 |
| 8. Sepsis | GSE5663 (Mild CLP Sepsis) | GSE9960 |
| 9. Sepsis | GSE5663 (Mild CLP Sepsis) | GSE28750 |
| 10. Sepsis | GSE5663 (Mild CLP Sepsis) | GSE13015 (GPL6947) |
| 11. Sepsis | GSE5663 (Mild CLP Sepsis) | GSE13015 (GPL6106) |
| 12. Sepsis | GSE5663 (CLP Sepsis) | GSE13904 |
| 13. Sepsis | GSE5663 (CLP Sepsis) | GSE9960 |
| 14. Sepsis | GSE5663 (CLP Sepsis) | GSE28750 |
| 15. Sepsis | GSE5663 (CLP Sepsis) | GSE13015 (GPL6947) |
| 16. Sepsis | GSE5663 (CLP Sepsis) | GSE13015 (GPL6106) |
| 17. Sepsis | GSE26472 (SPS2 Sepsis) | GSE13904 |
| 18. Sepsis | GSE26472 (SPS2 Sepsis) | GSE9960 |
| 19. Sepsis | GSE26472 (SPS2 Sepsis) | GSE28750 |
| 20. Sepsis | GSE26472 (SPS2 Sepsis) | GSE13015 (GPL6947) |
| 21. Sepsis | GSE26472 (SPS2 Sepsis) | GSE13015 (GPL6106) |
| 22. Sepsis | GSE19668 (AJ Strain, SA) | GSE13904 |
| 23. Sepsis | GSE19668 (AJ Strain, SA) | GSE9960 |
| 24. Sepsis | GSE19668 (AJ Strain, SA) | GSE28750 |
| 25. Sepsis | GSE19668 (AJ Strain, SA) | GSE13015 (GPL6947) |
| 26. Sepsis | GSE19668 (AJ Strain, SA) | GSE13015 (GPL6106) |
| 27. Sepsis | GSE19668 (C57 Strain, SA) | GSE13904 |
| 28. Sepsis | GSE19668 (C57 Strain, SA) | GSE9960 |
| 29. Sepsis | GSE19668 (C57 Strain, SA) | GSE28750 |
| 30. Sepsis | GSE19668 (C57 Strain, SA) | GSE13015 (GPL6947) |
| 31. Sepsis | GSE19668 (C57 Strain, SA) | GSE13015 (GPL6106) |
| 32. Sepsis | GSE19668 (C57 and AJ Strains, SA) | GSE13904 |
| 33. Sepsis | GSE19668 (C57 and AJ Strains, SA) | GSE9960 |
| 34. Sepsis | GSE19668 (C57 and AJ Strains, SA) | GSE28750 |
| 35. Sepsis | GSE19668 (C57 and AJ Strains, SA) | GSE13015 (GPL6947) |
| 36. Sepsis | GSE19668 (C57 and AJ Strains, SA) | GSE13015 (GPL6106) |

Baseline correspondence between each mouse model and human *in vivo* dataset was assessed by differential expression analysis and Gene Ontology (GO) pathway enrichment analysis of differentially expressed, homologous mouse and human transcripts (1). Differential expression was assessed by the Wilcoxon-Mann-Whitney (WMW) test with Benjamini Hochberg False Discovery Rate (FDR) correction (Significance: WMW p < 0.05, FDR q < 0.25). GO enrichment was performed on all DEGs using the Reactome pathway database annotation option in GO (21) (22, 23). We computed the precision and recall of the DEGs and pathways; precision represents the fraction of true positive results compared to all results, and recall represents the fraction of true positive results compared to all potentially positive results, with respect to correspondence between mouse and human datasets. These quantities are then summarized using two F-scores, which quantitatively characterize the precision and recall capabilities in single metrics – one for shared DEGs and one for shared enriched pathways. The F-score provided a summarized score that gave an equal weighting on both the accuracy of DEG and pathway predictions and how comprehensive the predictions were relative to the human-predicted associations.

The mouse model-predicted DEGs and enriched pathways constituted the baseline performance of mouse-to-human translation that our machine-learning approaches needed to improve upon to be considered successful (Figure 1A). We implemented supervised and semi-supervised versions of the k-nearest neighbors (KNN), support vector machine (SVM), random forest (RF), and artificial neural network (ANN) algorithms. In both the supervised and semi-supervised cases, we performed feature selection prior to training each classifier. We used Lasso or EN regularization as a feature selection method for the KNN, SVM, RF, and ANN classifiers. We varied the Lasso-EN regularization parameter α, using values of 1.0, 0.9, 0.7, 0.5, 0.3, and 0.1. In the case where α = 1, the feature selection method equivalent to Lasso and the smallest number of features were selected for classifier training. As α decreased, more features were selected to train the classifiers. We were thus able to assess the effect of the stringency of feature selection on model performance.

**Figure 1.**
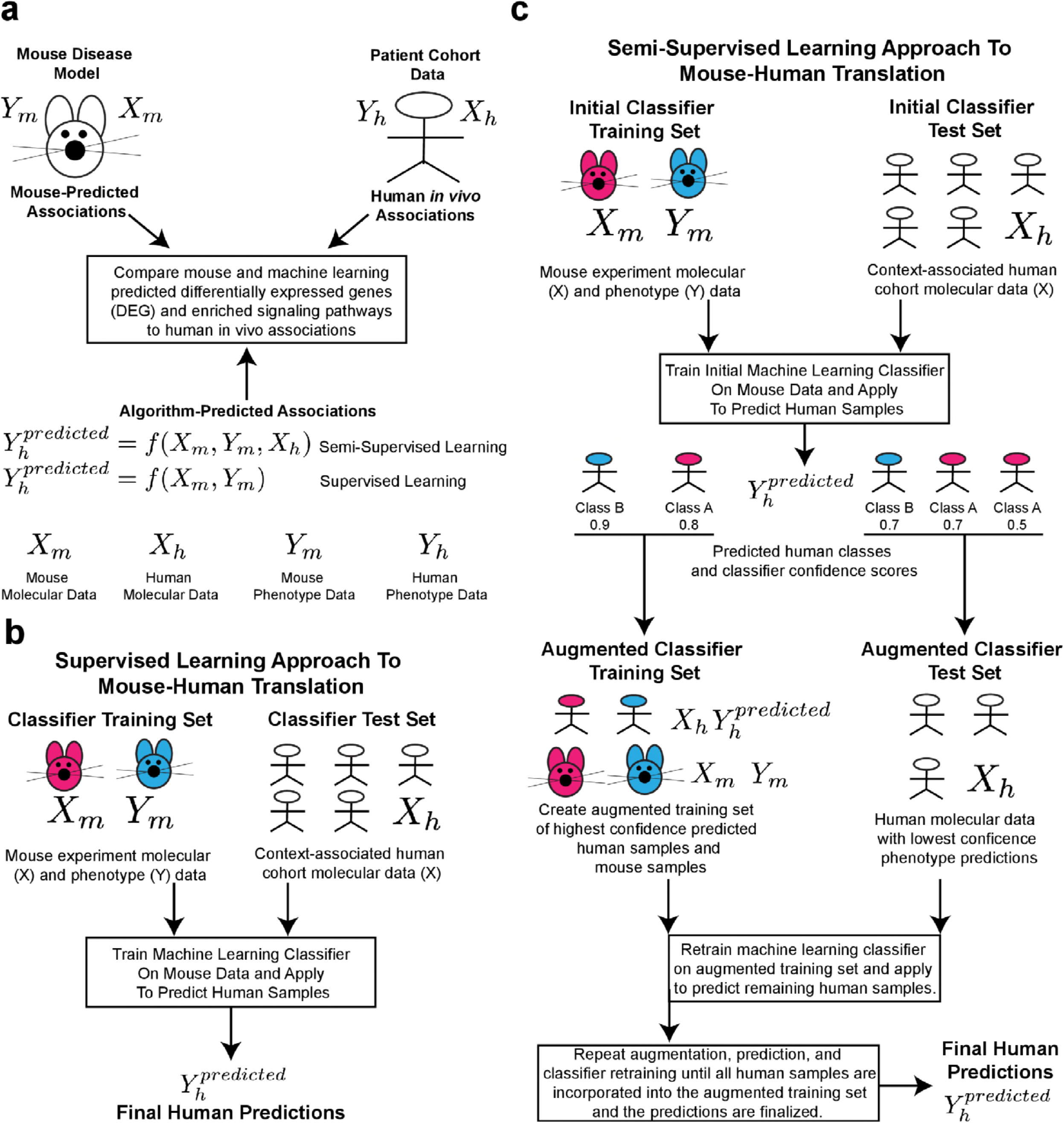
Conceptual framework for applying machine learning approaches to generalize molecular associations from mouse disease models to the human *in vivo* context. (a) Mouse model and machine learning-derived differentially expressed genes and enriched pathways are compared to human *in vivo* associations derived from patient gene expression datasets. (b) Supervised learning approaches utilize mouse data alone to construct a classifier that is then applied to human gene expression data to infer synthetic phenotype labels. The predicted human phenotypes are then used to derive predicted human *in vivo* DEGs and enriched signaling pathways. (c) Semi-supervised learning approaches begin by utilizing mouse model data to train an initial classifier to predict human synthetic phenotypes. Human samples with the highest confidence predicted synthetic phenotypes are then merged with the mouse cohort to create an augmented training dataset of mouse samples with measured phenotypes and human samples with simply predicted phenotypes. A new classifier is trained on this augmented mouse-human training set and applied to the remaining human samples. This procedure continues until all human samples have been merged with the training set at which point the classifications of the human samples are finalized. The predicted human phenotypes are then used to derive predicted human *in vivo* DEGs and enriched signaling pathways.

In the supervised case, the mouse molecule-phenotype dataset was used to train a machine learning classifier and the resulting classifier was applied to the human molecular dataset to infer synthetic phenotype labels (Figure 1B). In the semi-supervised case, the mouse dataset was first used to train a machine learning classifier in a supervised manner and then this classifier was applied to infer synthetic phenotype labels in the human dataset (Figure 1C). Then, following this first supervised training and prediction step, in a second step the human samples with the highest model confidence scores for their predicted phenotypes were merged with the mouse training dataset and used to construct a new classifier. The new classifier thus contained supervised information from the mouse dataset and unsupervised information from the human dataset. This procedure of algorithm training, prediction, and merging of the highest confidence predicted human samples with the training set continued until all human samples were merged with the training set. Once all human samples were merged with the training set, the performance of the machine learning classifier was assessed on the biological associations derived using the predicted human phenotypes. These associations were derived by differential expression and pathway enrichment analysis in the same manner as the human clinical datasets, with the predicted phenotypes used to define groups for comparison. These biological associations derived using the phenotypes predicted by the machine learning classifiers were summarized in DEG and pathway F-scores by comparing the algorithm-predicted DEGs and pathways to those from derived from the patient cohort data (Figure 1A).

### A semi-supervised neural network approach characterizes human *in vivo* molecule-phenotype relationships more accurately than mouse models alone

We assessed the performance of 1,728 machine learning classifiers (36 case studies × 6 a parameter values × 8 machine learning approaches) against the performance of the mouse-predicted DEG and pathway associations. The performance of each classifier was summarized by the area under the receiver operator characteristic curve (AUC) for the accuracy of the predicted human phenotypes and by the F-score of human DEGs and pathways inferred using predicted phenotypes. A generalized linear model (GLM) was trained to assess the impact of Lasso-EN regularization α values and the type of machine learning classifier on the AUC and DEG F-score performance metrics across all case studies. Neither the value of α (p = 0.725), nor the type of machine learning approach (p = 0.968) significantly impacted the AUC performance metric (Table S1). However, both α (p = 0.000242) and the type of machine learning method (p = 0.00189) significantly impacted the F-score metric (Table S2). The significance of the regularization parameter and classifier type for the F-score and not the AUC metric suggests that though each machine learning approach performed with comparable accuracy, the biological relevance of the predicted phenotypes were significantly influenced by the stringency of feature selection and choice of machine learning approach.

Since the F-score was a direct measure of the biological relevance of the predictions made by a particular algorithm, we focused on F-score as the most relevant performance metric. That is, we emphasized the capability for gaining biological insights over mere numerical predictive capacity. We computed the 95% confidence intervals of the F-scores for each machine learning approach and mouse model across all case studies and regularization parameters (Figure 2). The overall performance of mouse-derived DEGs for prospectively predicting human DEGs was surprisingly low (95% CI [0.090, 0.175], Average F-score = 0.132). For a machine learning approach to be considered successful, it needed to improve upon the performance of all mouse models across disease indications. Four machine learning approaches met this criteria. The semi-supervised ANN (ssANN), semi-supervised RF (ssRF), KNN, and SVM confidence intervals all excluded the mouse model F-score confidence interval indicating that these methods outperform the mouse model on average (Figure 2). However, the ssANN F-score confidence interval also excluded all other models indicating that the ssANN was the most successful model overall (95% CI [0.278, 0.375], Average F-score = 0.326). The ssANN similarly outperformed the mouse model and other computational approaches in terms of precision and recall (Table S3). We then performed WMW tests comparing the DEG F-scores between all pairs of machine learning approaches and confirmed that the ssANN performed significantly better than all other machine learning approaches (Table S4). Finally, we examined the average performance of the ssANN specifically across all case studies for each setting of the regularization parameter α (Table S5). An α value of 1.0 corresponding to Lasso regularization had the highest average F-score across all case studies (F = 0.381). Based upon the GLM, F-score confidence intervals, and average performance of the ssANN at each value of α, we concluded that the ssANN with Lasso regularization was the most broadly effective approach for prediction of human DEGs.

**Figure 2.**
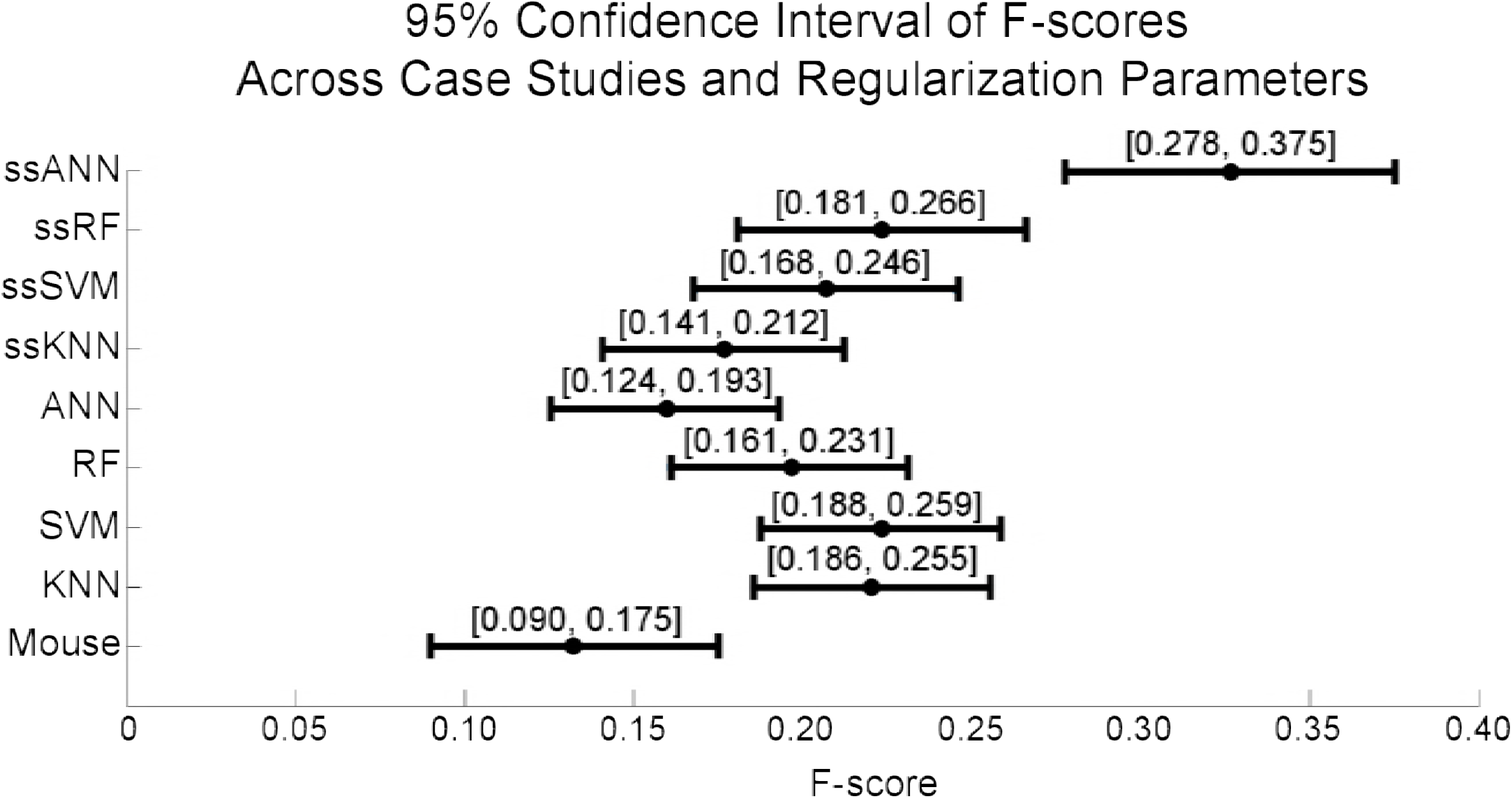
Identification of the most effective machine learning and regularization parameter. The 95% confidence intervals of the DEG F-scores of each machine learning approach across all regularization parameter values and case studies. The model confidence interval was computed from the DEG F-scores across all mouse-human case studies and the average F-score is denoted in the confidence interval.

### ssANNs better predict human *in vivo* genes and pathways than mouse models across disease contexts

Having identified the ssANN with Lasso regularization as the most broadly effective machine learning method for predicting human *in vivo* DEGs, we next compared the performance of the ssANN predictions of DEGs and enriched pathways to those made by mouse models in each individual case study (Figure 3, Table S6). In most cases, the mouse model pathway F-score is higher than the DEG F-score indicating that the mouse models considered here are more predictive of human *in vivo* pathway function than individual differential expression events (Figure 3a). Notably, the baseline correspondence between the enriched pathways identified by mouse models and human *in vivo* contexts was relatively consistent across disease indications, (Figure 3a), suggesting that mouse models of inflammatory pathologies recapitulate similar proportions of human *in vivo* molecular biology across indications independent of disease etiology complexity.

**Figure 3.**
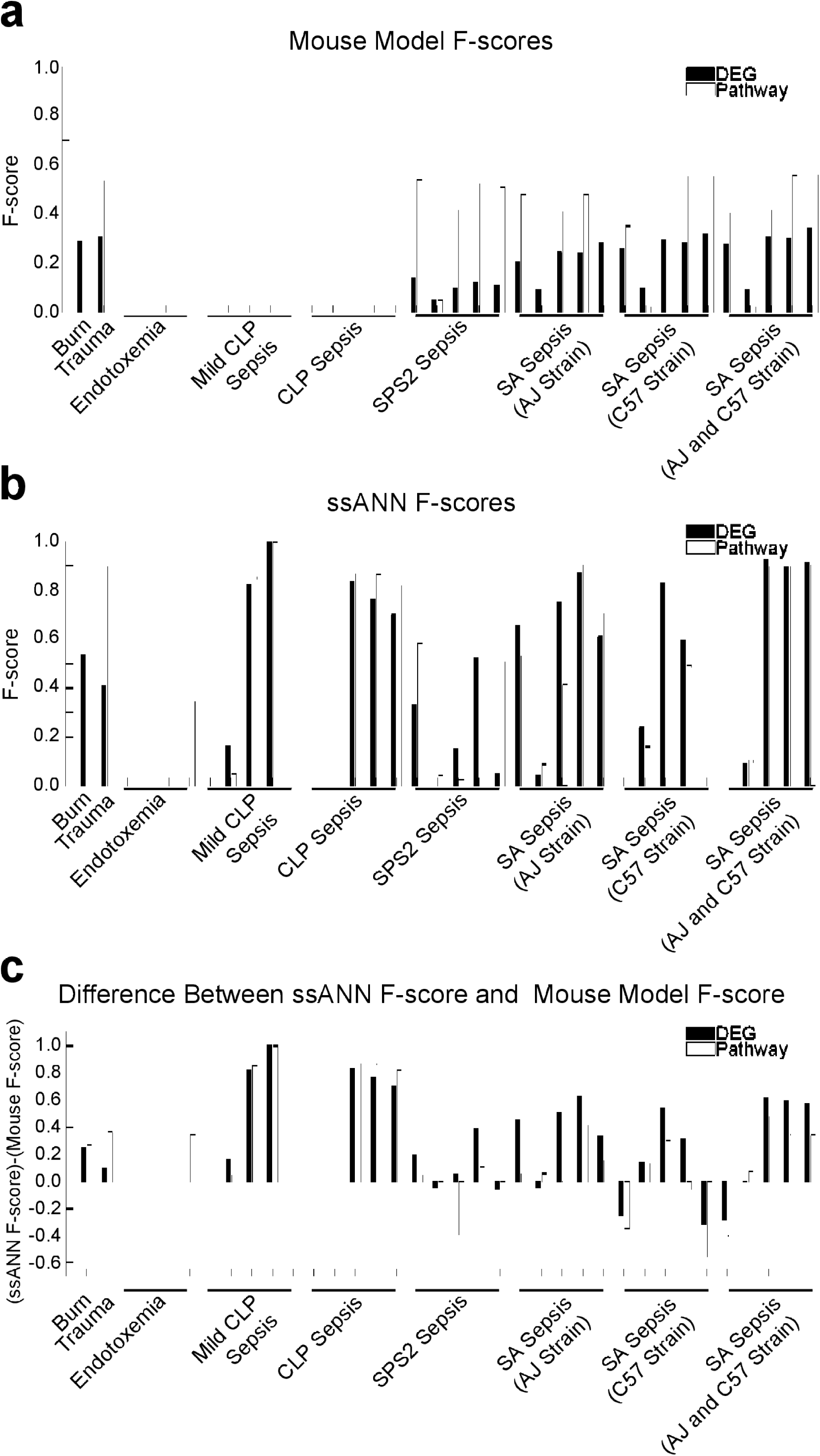
Comparison of mouse and ssANN DEG and pathway F-scores by individual disease context and case study. (a) Mouse model F-scores by case study. (b) Lasso regularized ssANN F-scores by case study. (c) Difference between ssANN and mouse model F-scores.

Notable exceptions to this pattern of mouse-human pathway correspondence were the endotoxemia and cecal ligation and puncture (CLP) mouse models. None of the endotoxemia or CLP driven sepsis mouse models, 14 of 36 case studies, had any DEGs at permissive statistical thresholds (WMW p < 0.05, FDR q < 0.25) (Figure 3a). This appears to contradict previous studies suggesting that CLP is the “gold standard” mouse model for mimicking human sepsis pathophysiology (24). Those previous studies focused on particular cytokines and higher order disease phenotype measures and did not subject the mouse model to the standard of prediction of human *in vivo* molecular changes as we do here.

In comparison, in 7 of these 14 CLP mouse model cases, the ssANN was able to correctly characterize a large proportion of human *in vivo* sepsis DEGs and pathways despite being trained on mouse models with no univariate differential expression events (Figure 3b). Here, the semi-supervised approach provides substantial benefit when mouse models, such as CLP-driven sepsis and LPS stimulated endotoxemia, do not recapitulate molecular features of human disease biology. Compared to the mouse models, the ssANN predicted an equal or greater proportion of human DEGs and enriched pathways in 29 of 36 case studies (Figure 3c). In the remaining 7 cases, the mouse models of *streptococcus pneumoniae serotype* 2 (SPS2) *and staphylococcus aureus* (SA) driven sepsis outperformed the ssANN in predicting DEGs and pathways in particular human cohorts. A single human dataset, GSE13015, a sepsis cohort where many of the patients had other infections such as melioidosis, was implicated in 4 of these 7 case studies (9). This suggests that the C57 strain mouse with an SA or SPS2-driven sepsis is an unusually satisfactory direct model for human sepsis with other infectious complications. The ssANN may have failed to outperform the mouse in these cases due to the heterogeneity of infections in the human cohort. This interpretation is supported by the fact that the ssANN outperforms the combined mouse cohort by a wide margin when the AJ and C57 mouse models are combined into a single training cohort (Figure 3c), suggesting that in a prospective applications to predict biological associations in a heterogeneous human cohort, the ssANN should be trained on a heterogeneous mouse cohort.

### Computational ssANN translation improves recall of human *in vivo* pathways in sepsis

This diversity of sepsis mouse models in our cohort made it possible to assess the correspondence of different protocols for generating sepsis mouse models to the human disease context. While CLP mouse models failed to identify any DEGs, the SPS2 and SA sepsis mouse models were both partially predictive of DEGs and pathways in human sepsis cohorts (Figure 3a). The SA mouse sepsis cohort was comprised of two different mouse strains, the highly susceptible A/J mouse strain and the somewhat resistant C57BL/6J strain (8). We were therefore able to compare four cohorts of sepsis models (SPS2- C57BL/6J, SA-A/J, SA-C57BL6J, and SA-mixed (A/J and C57BL6J)) in order to identify the most representative mouse models of clinical sepsis. We were also able to determine the DEGs and pathways of clinical sepsis implicated in particular mouse sepsis models recoverable by use of the ssANN.

Since enriched pathway predictions had a greater correspondence to human sepsis than DEGs alone, we compared the pathway associations derived from each sepsis mouse model to one another to identify common and distinguishing features of each model (Figure 4a). In total, 442 pathways were enriched across all human sepsis cohorts and multiple mouse sepsis models correctly predicted subsets of these pathways. All mouse models and strains correctly identified a set of 112 pathways including signaling by FGFR1, FGFR2, FGFR3, and FGFR4, and MAPK1 signaling (Table S7). This pathway signature of human sepsis appears to be highly reproducible in multiple mouse sepsis models, rendering it a stable signature for assessing therapeutic interventions and benchmarking mouse sepsis models against human data.

**Figure 4.**
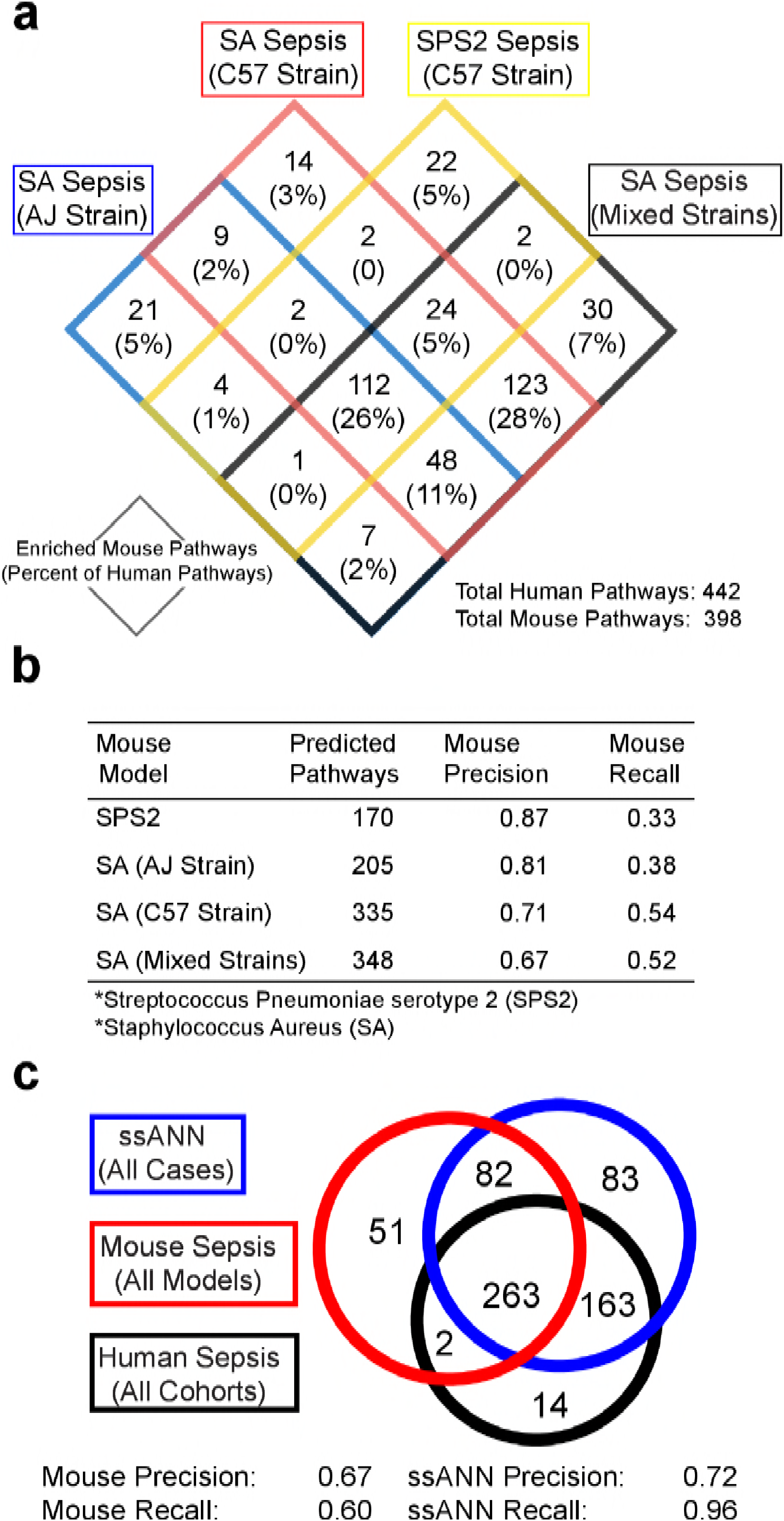
Assessing the correspondence of signaling pathways implicated by mouse models of sepsis or the ssANN to human *in vivo* sepsis. (a) Comparison of signaling pathways enriched in different strains and mouse models of sepsis and proportion of these pathways implicated by mouse models enriched in human sepsis *in vivo*. (b) Precision and recall of each mouse model’s predicted human *in vivo* signaling pathways by strain and type of mouse model. (c) Comparison of signaling pathway predictions by any sepsis mouse model, ssANN, and human *in vivo* sepsis.

We examined the precision and recall statistics of each mouse model across all human cohorts and observed that increasing the heterogeneity of the mouse cohort and aggregating predictions across multiple mouse models improves the coverage of human sepsis pathway predicted, but simultaneously degrades the precision of these predicted pathway signatures (Figure 4b). This contrasts with our finding that increasing the heterogeneity of the mouse cohort improved the predictive power of the ssANN (Figure 3c). Because heterogeneity of the mouse cohort degrades predictions of human pathways in the absence of the ssANN, this suggests that a heterogeneous mouse cohort contains latent features that the ssANN is able to detect and incorporate into its predictions of human *in vivo* pathways.

We then compared the combined pathway predictions of all mouse sepsis models to the predictions of the ssANN across all sepsis cases to assess correspondence with human *in vivo* sepsis pathway signatures (Figure 4c). The mouse sepsis models confirmed two pathways that the ssANN missed: the CD28-dependent VAV1 pathway and the oxidative stress induced senescence pathway. The oxidative stress senescence pathway was implicated by both of the SA mouse models in isolation, but not the mixed cohort, while the CD28-dependent VAV1 pathway was specifically implicated in the C57 strain. Use of a CD28 mimetic peptide has been shown to increase survival in gram-negative and polymicrobial models of mouse sepsis and is currently being explored as a therapeutic option for human sepsis (25). Though the mouse model recapitulated these two pathways missed by the ssANN, the ssANN performed with comparable precision to the mouse models overall (precision = 0.72) and recovered a strikingly higher proportion of *in vivo* human sepsis pathways (recall = 0.96) (Figure 4c). Furthermore, the ssANN recovered a set of 163 pathways enriched in human sepsis *in vivo* that were not identified in any mouse models of sepsis (Table S8). These pathways included thrombin signaling, TGFβ signaling, as well as several RNA transcriptional and post-translational modification-based pathways (Table S8) that all mouse models of sepsis lacked. Both thrombin and TGFβ signaling have been shown to play key roles in the pathology of sepsis and have been investigated for therapeutic and prognostic applications in sepsis (26, 27) (28). This result suggests that combining context-associated human data with mouse disease model data recovers important aspects of human *in vivo* signaling without sacrificing the baseline precision of the mouse.

## DISCUSSION

In this study we demonstrate that prospective translation of biological insights from mouse models to human inflammatory disease contexts was best accomplished by a semi-supervised artificial neural network. The ssANN correctly predicted human *in vivo* biological associations by combining supervised modeling of mouse model datasets with unsupervised modeling of patient molecular datasets. Though the supervised KNN, SVM, and semi-supervised RF were all superior to the mouse models overall, the ssANN outperformed each of these methods making it the most broadly effective approach for mouse to human insight translation across case studies (Figure 2). It is striking that the overall F-scores of mouse models for predicting human DEGs was merely 0.132, indicating that there is a substantial portion of human biology overlooked in established mouse models of inflammatory disease. However, application of a ssANN within the framework we have proposed here recovers 2.5x as many true human biological associations on average (F score = 0.326), improved upon the mouse in 29 of 36 case studies, and achieved human *in vivo* pathway prediction F-scores over 0.70 in 13 of 36 cases. Thus, systems modeling aided translation of insights from mouse to human contexts offers substantial benefits in characterizing human molecule-phenotype associations when human phenotype data for the indication of interest is not available.

The diversity of sepsis mouse model types in our case studies allowed us to assess the correspondence of different types of sepsis mouse models to human *in vivo* sepsis and compare the sepsis mouse models overall to the ssANN. We found that the CLP mouse model of sepsis is a poor predictor of human *in vivo* DEGs and enriched pathways. However, the SA and SPS mouse models share a pathway signature accounting for 26% of all human *in vivo* sepsis signaling (Figure 4a). Despite this, when all predictions of all sepsis mouse models are combined, the overall precision of the mouse models was 0.67 and the total coverage of human *in vivo* sepsis pathways 60%. The aggregated predictions of the ssANN improved upon this result by a striking margin, recovering 96% of human *in vivo* sepsis pathways with a precision of 0.72. This amounted to identification of 163 enriched pathways dysregulated in human *in vivo* sepsis that all mouse models of sepsis lacked (Table S8). By comparing the predicted enriched signaling pathways from each sepsis mouse model, we were able to assess the consistency of these models with one another as well as with human *in vivo* sepsis (Figure 4a). The diversity of predictions by these mouse models and overall low recall of each mouse model individually illustrates the challenge of mimicking human sepsis by means of mouse models where the causal mechanisms are single-agent perturbations (Figure 4b). Human sepsis is a considerably more complex system and though mouse models will likely continue to improve in fidelity, a wide gap in molecular correspondence between the two species remains.

The role of the heterogeneity of the mouse sepsis cohort differed between interpreting the mouse predictions directly and using the ssANN to predict human *in vivo* biology. When aggregated together, the mouse model predictions of human sepsis pathways increased in total coverage, but tended to be less precise. This runs contrary to observations that including multiple mouse strains in an experiment and using the increased heterogeneity of the mouse cohort as a surrogate for human disease heterogeneity improved translation of insights from mouse models to human context (29). In the context of the ssANN, increasing the heterogeneity of the mouse cohort in the training set by inclusion of multiple strains resulted in better predictions than when the ssANN was trained on individual mouse sepsis strains, particularly when the human cohort being predicted was itself heterogeneous (Figure 3b).

The principles of semi-supervised learning approaches share many features with those of transfer learning, the philosophy that combining information from multiple domains can improve the performance of machine learning approaches for extracting biological insights. In particular, transfer learning has been applied to problems of spatial localization of proteins in cells as well as to prediction of DNA splicing sites (30) (31). In these applications, a set of training data (X_train_ and Y_train_) are integrated with a context-related dataset (X_context_) to improve the performance of the algorithm in an approach known as inductive transfer learning. Our approach is an example of transductive transfer learning. In this case, X_context_ = X_test_ and the test dataset is incorporated into the algorithm training procedure in an unsupervised manner. The success of a transductive machine learning approach is determined by the robustness of the approach to different combinations of training and testing datasets. We assembled the literature case studies in this paper to address requirement, using all combinations of training and testing datasets for a disease indication and requiring a classification model to work in a variety of contexts to be judged a success.

The ideal case for characterizing the biology of human disorders would be the availability of comprehensive human phenotype and molecular data from clinical cohorts. However, since novel perturbations to the disease system cannot be studied in the human *in vivo* context, mouse disease model systems and emerging human *in* vitro model systems will continue to play an important role in biomedical research. It is in this context that we propose a delineation of four categories of *Translation Problems*, those of generalizing insights from model systems to human *in vivo* contexts (Figure 5). The most challenging case is when only model system molecular and phenotype data are available (Category 4). If human-based prior knowledge, such as candidate genes or clinical observations, is available to integrate with model system data, then generalization can be characterized as a Category 3 problem. In Category 2 problems, condition-specific human molecular data is available to combine with model system molecular and outcome data to characterize human biology. Inferences from solving Category 2 problems can be further refined with human-based prior knowledge to refine biological inferences in a Category 1 problem.

**Figure 5.**
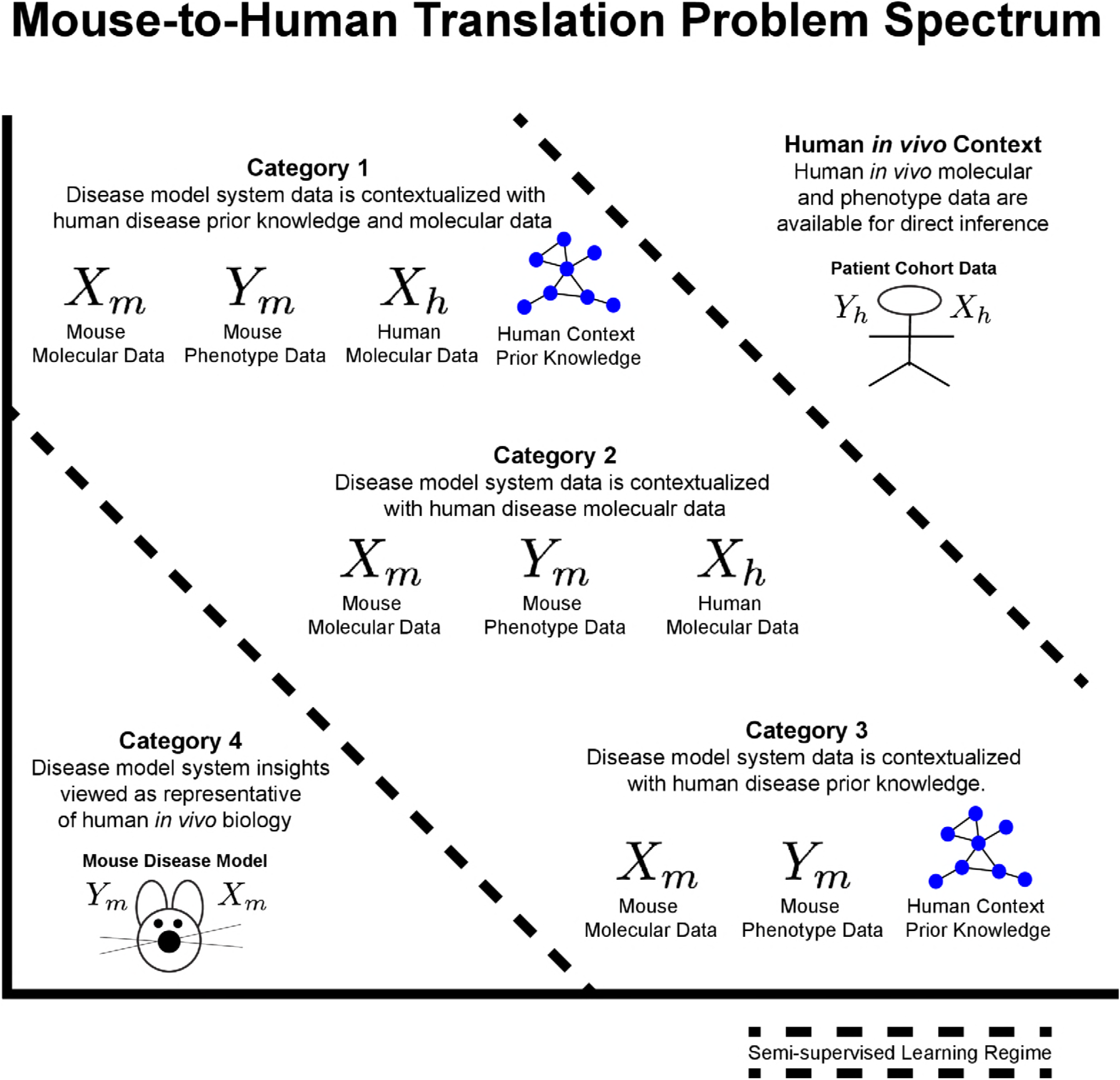
Delineation of four categories of Translation Problems, those of generalizing insights from model systems to human *in vivo* contexts. The type of datasets available determines the category, though Categories 1-3 are all potentially approachable by integration of model system data with human disease-context data using semi-supervised learning approaches.

We submit that the majority of previous biological research falls into Category 4 translation problems. Seok *et al.* illustrated the potential problems of direct generalization of mouse insights and of translational science following a Category 4 approach (1). By preselecting genes of interest based on known human biology, Takao *et al.* do not engage with the issues and challenges of prospectively translating insights from mouse to human contexts when information about human phenotypes is not available for the mouse study perturbation.

Within this framework, our efforts here are best viewed as a generalized solution to Category 2 translation problems. Another field where Category 2 translation problems frequently emerge is in pharmacogenomics studies where molecular features are connected to drug response outcomes in a model system and used as indicators of human *in vivo* mechanism. Many such experiments examine novel individual or novel combinations of compounds untested in patients illustrating the scarcity of human *in vivo* phenotype data. Thus, interpretation of these experimental results represents a large body of Category 2 translation problems where inferring human *in vivo* therapeutic biomarkers from nonhuman model system screening data is the objective. Our results here suggest that such experiments would be better interpreted if integrated with available context-associated human data using a semi-supervised ANN.

Despite advances in the fidelity of model system biology to human contexts, generalizability of findings of model system experiments will continue to be a key issue in both basic biology and translational science research (32, 33). Whenever the model system data alone forms the basis of inference, whether through direct interpretation or indirectly through a computational description of the model system’s biology, key aspects of human biology are likely to be overlooked or misrepresented. Transductive transfer learning approaches that neither aim for a generalizable computational model nor rely on the model system training data alone, recover more relevant human *in vivo* biology as a downstream consequence of creating good predictions of human phenotype for a specific patient cohort. This conceptual shift from direct interpretations of model system data to the indirect generalization of model system biology through integration with human data in semi-supervised learning framework has the potential to improve the success rate of translating model system therapies and insights to human patients.

## MATERIALS AND METHODS

### Dataset Collection and Processing

The datasets used in our analysis were obtained from Gene Expression Omnibus (34) and selected based on their inclusion in two papers published in the Proceedings of the National Academy of Science (PNAS) titled “Genomic Responses in Mouse Models Poorly Mimic Human Inflammatory Diseases” and “Genomic Responses in Mouse Models Greatly Mimic Human Inflammatory Diseases”(1, 2). Since we used the human datasets as test datasets and the mouse datasets as training datasets for machine learning applications, we applied the additional criteria that phenotypes and tissues of origin were comparable between mouse model and human *in vivo* datasets to ensure comparable training and test cases for algorithm performance comparison. Based on these criteria, we excluded the acute respiratory distress syndrome and acute infection datasets, and mouse splenocyte samples from GSE7404, GSE5663 antibiotic treated sepsis mice spleen samples, and GSE26472 mouse liver and lung samples.

The final cohort consisted of 6 mouse cohorts (3 sepsis datasets (8, 19, 20), 1 burn(18), 1 trauma(18), 2 endotoxemia(18, 19)) and 7 human cohorts (4 sepsis(9, 14-17), 1 burn(1, 10), 1 trauma(11), 1 endotoxemia(13)) (Table 1). Mouse probe identifiers were converted to gene symbols and mapped to homologous human genes using the mouse genome informatics database(35, 36). If multiple diseases or microarray platforms were used in a dataset, the dataset was partitioned by disease type to and array platform to create multiple case studies. All pairs of mouse and human datasets for a given indication constituted 36 case studies with comparable mouse and human tissues and phenotypes (Table 2). Duplicate genes in each dataset in each case study were removed by retaining only those genes with the maximum average expression across all samples. Datasets were z-scored by gene.

### Supervised and Semi-Supervised Classification Models

We implemented supervised and semi-supervised versions of the k-nearest neighbors (KNN), support vector machine (SVM), random forest (RF), and artificial neural network (ANN) algorithms. KNN was trained using three nearest neighbors. RF models were constructed from 50 decision trees. The ANN was a feed-forward neural network with three layers. The input layer consisted of one node for each feature, the output layer consisted of two nodes, one for each class, and the hidden layer consisted of the average of the number of input and output nodes rounded up to the nearest integer. ANN synapse weights were computed using scaled conjugate gradient backpropagation. All models were implemented in MATLAB 2016a.

Prior to training the algorithms, we performed feature selection with either Lasso or elastic net (EN) regularization. Different values of the regularization parameter α were examined in order to assess the impact of varying the number of features selected for training the supervised and semi-supervised classifiers (α = 1.0, 0.9, 0.7, 0.5, 0.3, 0.1). In the case of supervised classification models, Lasso and EN regularization underwent 10-fold cross validation (leave one out cross validation for mouse endotoxemia dataset GSE5663) to learn a set of features minimizing the mean squared error of the prediction of mouse phenotypes. These features were then used to train a supervised classifier (KNN, SVM, RF, or ANN) on the mouse dataset. The supervised classification model was then applied to the human dataset for that particular case study to infer synthetic human phenotypes (Figure 1B). In the case of semi-supervised classification models, Lasso or EN regularization was performed on the mouse dataset for a case study in the same manner as for supervised classification models. These features were then used to train an initial supervised classification model on the mouse data alone to predict the human samples’ synthetic phenotypes. Following this initial training and prediction step, the human samples with the highest 10% of confidence scores on their predicted phenotypes were combined with the mouse dataset to create a new augmented training set (Figure 1C). In the second iteration, feature selection and model training proceeded using this training set of mouse and human samples. All human samples in the test set were re-classified and the confidence score threshold of inclusion was dropped to 20%. The procedure of feature selection, model retraining, classification, and training set augmentation by lowering the score threshold 10% continued until all human samples are incorporated into the training set. Since ANN training is inherently stochastic, we specified that the semi-supervised ANN would proceed to the second iteration only if more than one human sample was classified into each class. If this condition as not met after 50 training iterations, the semi-supervised ANN proceeded with further training and prediction iterations on the human dataset using an initial model that did not have human predicted phenotypes in both classes.

### Model Performance Assessment

Classification models were evaluated by their ability to discriminate between human phenotypes and by the extent to which analyzing the human molecular data using the predicted human phenotypes implicated the same genes as using the true human phenotypes. Classification performance was assessed by the area under the receiver operating characteristic curve (AUC) for the test set of human samples.

Differential expression analysis was performed on the homologous mouse and human genes in each case study using the phenotypes from the original datasets to identify differentially expressed mouse and human genes. Following application of a classification model, differential expression analysis was then performed on the human dataset using the predicted synthetic phenotypes generated by the algorithm. Differential expression was assessed by the Wilcoxon-Mann-Whitney (WMW) test with Benjamini Hochberg False Discovery Rate (FDR) correction. This nonparametric differential expression analysis approach was necessary because previous studies have shown that these mouse and human inflammatory pathology datasets do not satisfy the statistical assumptions required for parametric model-based differential expression analysis (2). Genes were considered differentially expressed if both WMW p < 0.05 and FDR q < 0.25. GO enrichment was performed on all DEGs in each case study, for the human patient data, mouse data, and patient data with predicted synthetic phenotypes using the Reactome pathway database annotation option in GO (21) (22, 23).

A DEG or enriched pathway identified in the mouse model was considered a true positive (TP) if that gene or pathway was implicated in the human clinical gene expression data. False negatives (FN) were DEGs or enriched pathways implicated in the human data, but not implicated by the mouse model. False positives (FP) were DEGs or pathways implicated in the mouse but not in the human data. DEGs and pathways identified using the synthetic human phenotypes generated machine learning approaches were considered TP, FP, and FN by their correspondence to the DEGs and pathways implicated in human clinical gene expression data. We computed the precision and recall for the DEGs predicted by the mouse model and machine learning classifiers and aggregated these into an F-score for each prediction modality:

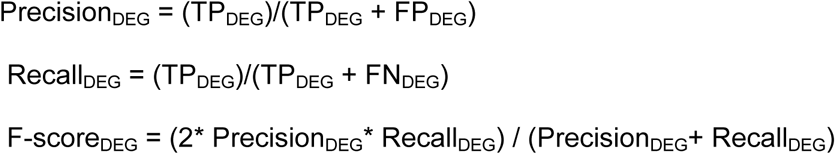

Enriched pathway precision, recall, and F-scores were analogously computed for TP, FP, and FN predicted pathways from the mouse model and machine learning classifiers:

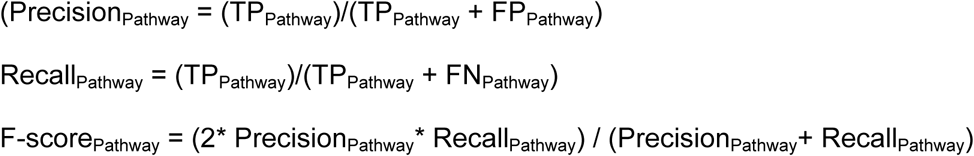

### Code Availability

All models and analyses were implemented in MATLAB 2016b. KNN, SVM, and RF functions were implemented using the fitcknn, fitcsvm, and TreeBagger functions respectively. Neural networks were implemented using the MATLAB Neural Network Toolbox. Semi-supervised learning algorithms are available as a supplementary file.

## LIST OF SUPPLEMENTARY MATERIAL

**Table S1.** Generalized linear model coefficients for the effect of model type, and elastic net parameter on the AUC performance of machine learning classifiers.

**Table S2.** Generalized linear model coefficients for the effect of model type, and elastic net parameter on the F-score performance of machine learning classifiers.

**Table S3.** Precision and recall confidence intervals across all case studies and regularization parameters.

**Table S4.** Wilcoxon Mann Whitney tests comparing the F-scores for each machine learning approach to identify the most effective method.

**Table S5.** Average F-score across all case studies by choice of machine learning method and regularization parameter alpha.

**Table S6.** Summary of the number of DEGs and enriched pathways associated with each human patient cohort, mouse model cohort, and ssANN predicted associations.

**Table S7.** Reactome pathways enriched in human sepsis in vivo and consistently recovered by the SA and SPS2 sepsis mouse models.

**Table S8.** Reactome pathways enriched in human sepsis in vivo, missed by both the SA and SPS2 sepsis mouse models, but recovered by the ssANN.

## AUTHOR CONTRIBUTIONS

DKB performed analyses, designed the study, and wrote the manuscript. EAP, KMH, and DAL designed the study and wrote the manuscript.

## ACKNOWLEDGEMENTS

The authors wish to thank John Hambor, Erick Young, Kevin Janes, Brian Joughin, Alina Starchenko, Evan Chiswick, and Samanta Dale Strasser for their helpful commentary.

## REFERENCES

1. Seok J, Warren HS, Cuenca AG, Mindrinos MN, Baker HV, Xu W, et al. Genomic responses in mouse models poorly mimic human inflammatory diseases. Proc Natl Acad Sci U S A. 2013;110(9):3507–12.

2. Takao K, Miyakawa T. Genomic responses in mouse models greatly mimic human inflammatory diseases. Proc Natl Acad Sci U S A. 2015;112(4):1167–72.

3. Domcke S, Sinha R, Levine DA, Sander C, Schultz N. Evaluating cell lines as tumour models by comparison of genomic profiles. Nat Commun. 2013;4:2126.

4. Goodspeed A, Heiser LM, Gray JW, Costello JC. Tumor-Derived Cell Lines as Molecular Models of Cancer Pharmacogenomics. Mol Cancer Res. 2016;14(1):3–13.

5. Jiang G, Zhang S, Yazdanparast A, Li M, Pawar AV, Liu Y, et al. Comprehensive comparison of molecular portraits between cell lines and tumors in breast cancer. BMC Genomics. 2016;17 Suppl 7:525.

6. Nickerson ML, Witte N, Im KM, Turan S, Owens C, Misner K, et al. Molecular analysis of urothelial cancer cell lines for modeling tumor biology and drug response. Oncogene. 2017;36(1):35–46.

7. Kodamullil AT, Iyappan A, Karki R, Madan S, Younesi E, Hofmann-Apitius M. Of Mice and Men: Comparative Analysis of Neuro-Inflammatory Mechanisms in Human and Mouse Using Cause-and-Effect Models. J Alzheimers Dis. 2017;59(3):1045–55.

8. Ahn SH, Deshmukh H, Johnson N, Cowell LG, Rude TH, Scott WK, et al. Two genes on A/J chromosome 18 are associated with susceptibility to Staphylococcus aureus infection by combined microarray and QTL analyses. PLoS Pathog. 2010;6(9):e1001088.

9. Pankla R, Buddhisa S, Berry M, Blankenship DM, Bancroft GJ, Banchereau J, et al. Genomic transcriptional profiling identifies a candidate blood biomarker signature for the diagnosis of septicemic melioidosis. Genome Biol. 2009;10(11):R127.

10. Peterson JR, De La Rosa S, Eboda O, Cilwa KE, Agarwal S, Buchman SR, et al. Treatment of heterotopic ossification through remote ATP hydrolysis. Sci Transl Med. 2014;6(255):255ra132.

11. Xiao W, Mindrinos MN, Seok J, Cuschieri J, Cuenca AG, Gao H, et al. A genomic storm in critically injured humans. J Exp Med. 2011;208(13):2581–90.

12. Foteinou PT, Calvano SE, Lowry SF, Androulakis IP. Multiscale model for the assessment of autonomic dysfunction in human endotoxemia. Physiol Genomics. 2010;42(1):5–19.

13. Calvano SE, Xiao W, Richards DR, Felciano RM, Baker HV, Cho RJ, et al. A network-based analysis of systemic inflammation in humans. Nature. 2005;437(7061):1032–7.

14. Wong HR, Cvijanovich N, Allen GL, Lin R, Anas N, Meyer K, et al. Genomic expression profiling across the pediatric systemic inflammatory response syndrome, sepsis, and septic shock spectrum. Crit Care Med. 2009;37(5):1558–66.

15. Sutherland A, Thomas M, Brandon RA, Brandon RB, Lipman J, Tang B, et al. Development and validation of a novel molecular biomarker diagnostic test for the early detection of sepsis. Crit Care. 2011;15(3):R149.

16. Tang BM, McLean AS, Dawes IW, Huang SJ, Lin RC. Gene-expression profiling of peripheral blood mononuclear cells in sepsis. Crit Care Med. 2009;37(3):882–8.

17. Payen D, Lukaszewicz AC. Gene-expression profiling of peripheral blood mononuclear cells in sepsis. Crit Care Med. 2009;37(7):2323–4; author reply 4.

18. Lederer JA, Brownstein BH, Lopez MC, Macmillan S, Delisle AJ, Macconmara MP, et al. Comparison of longitudinal leukocyte gene expression after burn injury or trauma-hemorrhage in mice. Physiol Genomics. 2008;32(3):299–310.

19. Chung TP, Laramie JM, Meyer DJ, Downey T, Tam LH, Ding H, et al. Molecular diagnostics in sepsis: from bedside to bench. J Am Coll Surg. 2006;203(5):585–98.

20. Weber M, Lambeck S, Ding N, Henken S, Kohl M, Deigner HP, et al. Hepatic induction of cholesterol biosynthesis reflects a remote adaptive response to pneumococcal pneumonia. FASEB J. 2012;26(6):2424–36.

21. Croft D, Mundo AF, Haw R, Milacic M, Weiser J, Wu G, et al. The Reactome pathway knowledgebase. Nucleic Acids Res. 2014;42 (Database issue):D472–7.

22. Ashburner M, Ball CA, Blake JA, Botstein D, Butler H, Cherry JM, et al. Gene ontology: tool for the unification of biology. The Gene Ontology Consortium. Nat Genet. 2000;25(1):25–9.

23. Gene Ontology C. Gene Ontology Consortium: going forward. Nucleic Acids Res. 2015;43 (Database issue):D1049–56.

24. Dejager L, Pinheiro I, Dejonckheere E, Libert C. Cecal ligation and puncture: the gold standard model for polymicrobial sepsis? Trends Microbiol. 2011;19(4):198–208.

25. Ramachandran G, Kaempfer R, Chung CS, Shirvan A, Chahin AB, Palardy JE, et al. CD28 homodimer interface mimetic peptide acts as a preventive and therapeutic agent in models of severe bacterial sepsis and gram-negative bacterial peritonitis. J Infect Dis. 2015;211(6):995–1003.

26. Bae JS, Lee W, Nam JO, Kim JE, Kim SW, Kim IS. Transforming growth factor beta-induced protein promotes severe vascular inflammatory responses. Am J Respir Crit Care M ed. 2014;189(7):779–86.

27. Ahmad S, Choudhry MA, Shankar R, Sayeed MM. Transforming growth factor-beta negatively modulates T-cell responses in sepsis. FEBS Lett. 1997;402(2-3):213–8.

28. Petros S, Kliem P, Siegemund T, Siegemund R. Thrombin generation in severe sepsis. Thromb Res. 2012;129(6):797–800.

29. Buscher K, Ehinger E, Gupta P, Pramod AB, Wolf D, Tweet G, et al. Natural variation of macrophage activation as disease-relevant phenotype predictive of inflammation and cancer survival. Nat Commun. 2017;8:16041.

30. Widmer C, Toussaint NC, Altun Y, Ratsch G. Inferring latent task structure for Multitask Learning by Multiple Kernel Learning. BMC Bioinformatics. 2010;11 Suppl 8:S5.

31. Breckels LM, Holden SB, Wojnar D, Mulvey CM, Christoforou A, Groen A, et al. Learning from Heterogeneous Data Sources: An Application in Spatial Proteomics. PLoS Comput Biol. 2016;12(5):e1004920.

32. Huh D, Matthews BD, Mammoto A, Montoya-Zavala M, Hsin HY, Ingber DE. Reconstituting organ-level lung functions on a chip. Science. 2010;328(5986):1662–8.

33. Domansky K, Inman W, Serdy J, Dash A, Lim MH, Griffith LG. Perfused multiwell plate for 3D liver tissue engineering. Lab Chip. 2010;10(1):51–8.

34. Edgar R, Domrachev M, Lash AE. Gene Expression Omnibus: NCBI gene expression and hybridization array data repository. Nucleic Acids Res. 2002;30(1):207–10.

35. Blake JA, Eppig JT, Kadin JA, Richardson JE, Smith CL, Bult CJ, et al. Mouse Genome Database (MGD)-2017: community knowledge resource for the laboratory mouse. Nucleic Acids Res. 2017;45(D1):D723–D9.

36. Eppig JT, Richardson JE, Kadin JA, Ringwald M, Blake JA, Bult CJ. Mouse Genome Informatics (MGI): reflecting on 25 years. Mamm Genome. 2015;26(7-8):272–84.

